# Altered BOLD signal variation in Alzheimer’s disease and frontotemporal dementia

**DOI:** 10.1101/455683

**Authors:** Timo Tuovinen, Janne Kananen, Riikka Rytty, Virpi Moilanen, Ahmed Abou Elseoud, Anne M Remes, Vesa Kiviniemi, ADNI

## Abstract

Recently discovered glymphatic brain clearance mechanisms utilizing physiological pulsations have been shown to fail at removing waste materials such as amyloid and tau plaques in neurodegenerative diseases. Since cardiovascular pulsations are a main driving force of the clearance, this research investigates if commonly available blood oxygen level-dependent (BOLD) signals at 1.5 and 3 T could detect abnormal physiological pulsations in neurodegenerative diseases. Coefficient of variation in BOLD signal (CV_BOLD_) was used to estimate contribution of physiological signals in Alzheimer’s disease (AD) and behavioural variant frontotemporal dementia (bvFTD). 17 AD patients and 18 bvFTD patients were compared to 24 control subjects imaged with a 1.5 T setup from a local institute. AD results were further verified with 3 T data from the Alzheimer’s disease neuroimaging initiative (ADNI) repository with 30 AD patients and 40 matched controls. Effect of motion and gray matter atrophy was evaluated and receiver operating characteristic (ROC) analyses was performed.

The CV_BOLD_ was higher in both AD and bvFTD groups compared to controls (p < 0.0005). The difference was not explained by head motion or gray matter atrophy. In AD patients, the CV_BOLD_ alterations were localized in overlapping structures in both 1.5 T and 3 T data. Localization of the CV_BOLD_ alterations was different in AD than in bvFTD. Areas where CV_BOLD_ is higher in patient groups than in control group involved periventricular white matter, basal ganglia and multiple cortical structures. Notably, a robust difference between AD and bvFTD groups was found in the CV_BOLD_ of frontal poles. In the analysis of diagnostic accuracy, the CV_BOLD_ metrics area under the ROC for detecting disease ranged 0.85 – 0.96.

**Conclusions:** The analysis of brain physiological pulsations measured using CV_BOLD_ reveals disease-specific alterations in both AD and bvFTD.

## Introduction

The two most common forms of early-onset dementia are Alzheimer’s disease (AD) and behavioral variant frontotemporal dementia (bvFTD). Multiple resting-state functional MRI (rs-fMRI) studies concerning AD and bvFTD have been published. In AD the findings have been relatively consistent, with reduced default mode network (DMN) connectivity reported in numerous studies, and it seems to correlate with disease severity [Agosta et al., 2012; Binnewijzend et al., 2012; Greicius et al., 2004; Hafkemeijer et al., 2012; Li et al., 2002; Zhou et al., 2010]. In bvFTD, reduced salience network (SLN) connectivity has been most reported finding, although there has been some diverseness [Farb et al., 2013; Filippi et al., 2013; Lee et al., 2014; Rytty et al., 2013; Zhou et al., 2010]. Advanced artifact removal affects the reproducibility of the functional connectivity findings [Griffanti et al., 2015; Tuovinen et al., 2017].

The blood oxygen level-dependent (BOLD) signal used in fMRI is an indirect marker of neuronal activity and also reflects other vascular, respiratory and physiological factors [Birn et al., 2006; Shmueli et al., 2007; Liu et al., 2013; Mark et al., 2015; Kiviniemi et al., 2016]. Brain activity induces a complex combination of changes in cerebral blood volume, cerebral blood flow, and oxygen extraction fraction that all affect the de-phasing of the water protons and modulate the T2*-weighted MRI signal intensity in the brain measured as BOLD signal [Buxton, 2012]. BOLD signal has marked correlation with cardiorespiratory pulsations due to the high sensitivity to blood flow status [Hoge et al., 1999; Wise et al., 2004; Birn et al., 2006; Chang et al., 2009; Chang and Glover, 2010; Birn et al., 2014]. Physiological fluctuations in BOLD signal have traditionally been considered as a nuisance concealing neural activity [Keilholz et al., 2017]

Physiological pulsations has been shown to be of vital importance for the homeostasis of the brain [Aspelund et al., 2015; Buxton, 2012; Buxton et al., 2014; Dreha-Kulaczewski et al., 2015; Erdő et al., 2017; Fleisher et al., 2009; Garrett et al., 2017; Glomb et al., 2018; Grady and Garrett, 2018; Iliff et al., 2013; Iliff et al., 2015; Martin et al., 2012; Nedergaard, 2013; Plog and Nedergaard, 2018]. The brain clearance driven by physiological pulsations has recently been strongly linked to neurodegenerative diseases [Iliff et al., 2012; Iliff et al., 2013; Iliff et al., 2014; Iliff et al., 2015; Kiviniemi et al., 2016; Kress et al., 2014; de Leon et al., 2017; Louveau et al., 2016; Peng et al., 2016; Plog et al., 2015; Snyder et al., 2015; Tarasoff-Conway et al., 2015]. The disease process may also alter the physiological noise structure or variability of the BOLD signal in the way it alters the low-frequency connectivity. There may be yet unknown physiological factors that have been overlooked in prior rs-fMRI data.

The temporal signal-to-noise ratio (tSNR) has been used to measure BOLD signal stability [Triantafyllou et al., 2005]. The inverse of tSNR, i.e., the temporal coefficient of variation of the BOLD signal (CV_BOLD_), is a similar quality assurance metric that also enables the detection of subtle artifacts from the fMRI data [Tuovinen et al., 2017]. Recently, CV_BOLD_ has been used to analyze changes in noise characteristics of BOLD data [Jahanian et al., 2014]. Using a similar approach, increased physiological fluctuations in white matter (WM) have been detected in AD [Makedonov et al., 2016] and small vessel disease [Makedonov et al., 2013]. CV_BOLD_ correlates with cerebral blood volume and cerebral blood flow measured using dynamic susceptibility contrast MRI in patients with acute ischemic stroke [Khalil et al., 2017].

Based on previous findings of abnormal noise characteristics of the BOLD data, it was hypothesized that the noise structure of the BOLD signal measured using CV_BOLD_ is altered in AD and bvFTD patients in specific ways. AD and bvFTD patients as well as healthy controls were imaged with a 1.5 T setup from a local institute. Results of AD patients were further verified with 3 T data from the Alzheimer’s disease neuroimaging initiative (ADNI) repository. Effect of motion and atrophy was evaluated.

## Materials and Methods

### Participants

The ethic committee of the Oulu University Hospital approved the study. Each participating site’s institutional review board approved the research protocols. Written informed consent was obtained from all participants or their legal guardians according to the Declaration of Helsinki.

### Local Institute Data

The study sample consisted of 17 AD patients, 18 bvFTD patients and 24 control subjects. The patients were examined at Oulu University Hospital at the Memory Outpatient Clinic of the Department of Neurology. They all underwent a thorough neurological and neuropsychological examination, screening laboratory tests and brain MRI that are routine in the clinic. All patients in the AD group met the NINCDS-ADRDA (National Institute of Neurological and Communicative Disorders and Stroke and the Alzheimer's Disease and Related Disorders Association) criteria for probable AD [McKhann et al., 1984]. Cerebrospinal fluid (CSF) AD biomarkers supported the diagnosis in all the cases with available results (n=12).

The bvFTD patients were clinically diagnosed according to the Neary criteria [Neary et al., 1998; Rascovsky et al., 2011]. Patients with progressive aphasia or semantic dementia phenotypes were excluded from the study. DNA samples were available for ten patients, and the *C9ORF72* repeat expansion was found in seven of them [Renton et al., 2011]. Mutations in progranulin or microtubule-associated protein tau genes were not found in any of the genetically tested bvFTD patients.

The control subjects were interviewed, and Mini-Mental State Examination (MMSE) and Beck’s Depression Inventory (BDI) were performed. No psychiatric or neurological disorders or medications affecting the central nervous system were allowed in the control group. Structural MRIs were interpreted as normal by clinical neuroradiologist.

The fMRI scan was performed within six months of the examination. The patients were allowed to continue their ongoing medications. Functional connectivity findings have been previously reported [Tuovinen et al., 2017]. These participants passed strict quality control prerequisites using methods published in that article.

### ADNI Data

To verify the main results from local institute data a reference dataset was obtained for AD and control groups. Data used in the preparation of this article were obtained from the ADNI database (http://adni.loni.usc.edu). The ADNI was launched in 2003 as a public-private partnership, led by Principal Investigator, Michael W. Weiner, MD. The primary goal of ADNI has been to test whether serial MRI, positron emission tomography (PET), other biological markers, and clinical and neuropsychological assessment can be combined to measure the progression of mild cognitive impairment (MCI) and early AD. For up-to-date information, see http://www.adni-info.org.

ADNI-2 participants with both resting-state fMRI and preprocessed anatomical scans within the first year of participation in the study were selected to minimize potential retention bias from repeat scans. Participants were included if they were between 55 and 82 years old, spoke English or Spanish as their first language, and had completed at least six years of schooling. The diagnostic classification was made by ADNI investigators using established criteria [McKhann et al., 1984]. Participants in the AD cohort fulfilled the NINCDS-ADRDA criteria for probable AD. Control subjects had MMSE scores between 24 and 30 and no significant memory concerns.

### Image Acquisition

#### Local Institute Data (1.5 T)

The subjects were imaged with a GE Signa HDx 1.5 T whole-body system with an eight-channel receiver coil. Subjects were given earplugs to reduce noise, and soft pads were fitted over the ears to protect hearing and to minimize motion. During MRI scanning all the subjects received identical instructions: to simply rest and focus on a cross on an fMRI-dedicated screen, which they saw through the mirror system of the head coil.

Structural high-resolution T1-weighted 3D FSPGR BRAVO images were taken under the following conditions: repetition time (TR) 12.1 ms, echo time (TE) 5.2 ms, flip angle (FA) of 20°, slice thickness 1.0 mm, field of view (FOV) 24.0 cm, matrix size 256 x 256 (i.e., 1 mm^3^ cubic voxels).

Resting-state BOLD data were acquired using a conventional gradient recalled echo-planar images (EPI) sequence under the following conditions: TR of 1800 ms, TE of 40 ms, 202 volumes (6 min 4 s), FA of 90°, 28 oblique axial slices, slice thickness 4 mm, inter-slice space 0.4 mm, covering the whole brain, FOV 25.6 cm x 25.6 cm, matrix size 64 x 64. The first three volumes were excluded from the time series due to T1 relaxation effects.

#### ADNI Data (3 T)

MRI images were collected on Philips 3 T MRI systems (Philips, Amsterdam, The Netherlands) from a total of 13 sites using a standardized protocol (http://adni.loni.usc.edu).

Structural T1 images were acquired with a TE of 3 ms, TR of 7 ms, FA of 9°, slice thickness of 1.2 mm, and a matrix size of 256 x 256 x 170. Preprocessing of the structural T1 images involved bias field correction using a histogram peak-sharpening algorithm (N3) [Sled et al., 1998] and was done already for the datasets downloaded.

Functional EPI images were acquired with a TR of 3,000 ms, TE of 30 ms, 140 volumes (7 min), FA of 80°, slice thickness of 3.3 mm, and a matrix size of 64 x 64 x 48. The first three volumes were excluded from the time series due to T1 relaxation effects.

#### Data Preprocessing

The BOLD rs-fMRI data were preprocessed with a typical FSL pipeline (http://www.fmrib.ox.ac.uk/fsl, FSL 5.0.8) including: head motion correction (FSL 5.0.8 MCFLIRT, motion estimates were also used in evaluating motion differences between groups), brain extraction (f = 0.5 and g = 0), spatial smoothing (Gaussian kernel 5-mm full width at half maximum), and high-pass temporal filtering by using a cutoff of 100 seconds. Multi-resolution affine co-registration within FSL FLIRT software was used to co-register mean, non-smoothed fMRI volumes to 3D FSGR volumes of corresponding subjects, and to co-register anatomical volumes to the Montreal Neurological Institute’s (MNI152) standard space template.

#### CV_BOLD_ maps

CV was used as a metric for the variation of fluctuations in the BOLD signal. The same method has been used in a study by [Jahanian et al., 2014] and is similar to the method used by [Makedonov et al., 2013; Makedonov et al., 2016].

For each preprocessed 4D fMRI dataset, a CV_BOLD_ map was calculated voxel-wise:
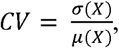

where X is voxel time series, *σ* is standard deviation and *μ* is mean.

Calculations were done using Matlab (version R2014b). Representative maps from one AD patient, one bvFTD patient and one control subject, as well as the group mean CV_BOLD_ maps, are shown in Fig. 1 for both local institute and ADNI data.

**Fig. 1.**
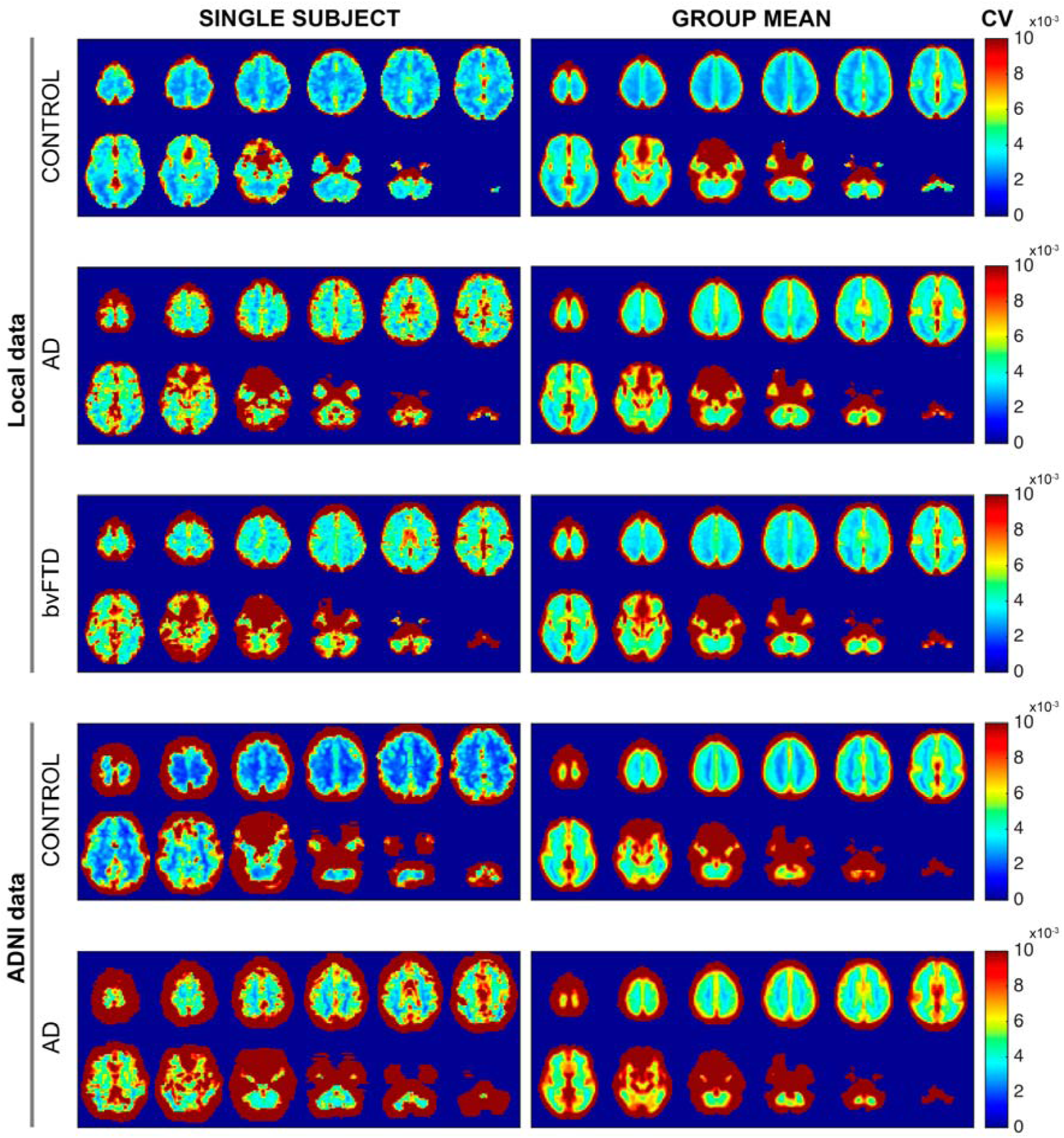
Coefficient of variation mapping of BOLD signal. Examples of randomly selected single subject and group mean CV_BOLD_ maps for both local institute and ADNI data in MNI152 space.

#### Regions-of-Interest analysis based on anatomical templates

ICBM152 nonlinear asymmetric 2009c [Fonov et al., 2009; Fonov et al., 2011] atlases (probabilistic map, thresholded to 50–100 %) were used as gray matter (GM), WM and CSF templates for the regions-of-interest (ROI) analysis. This approach was used in a study by [Jahanian et al., 2014] where they also showed that results are somewhat independent of the precise GM, WM and CSF segmentation strategy employed. From these ROIs, mean CV_BOLD_ was calculated subject-wise.

#### Voxel-level statistical analysis of CV_BOLD_ maps

Differences between study groups in the CV_BOLD_ maps were statistically tested using permutation-based nonparametric testing incorporating threshold-free cluster enhancement (TFCE) implemented in the FSL randomise tool with 10,000 random permutations [Smith and Nichols, 2009]. Resulting statistical maps were thresholded at p<0.05, 0.005 and 0.0005. The effect of GM atrophy on the CV_BOLD_ maps was also evaluated repeating the randomize analysis using the GM volume as a regressor. The resulting statistic maps were spatially correlated to the ones without a GM regressor using the fslcc tool from FSL.

#### Effect of motion

Motion estimates computed by the MCFLIRT algorithm in the preprocessing step was used to assess the effect of motion on the CV_BOLD_ values. Subject-wise absolute displacement vectors (in mm) were extracted, which describes the amount of movement in all directions over the whole scan as a marker of gross motion. Also, relative displacement vectors were extracted, as a marker of motion between each EPI volume. Both vectors were also averaged across volumes to get mean values. Additionally, maximum motion value and the number of peaks in the subject-wise motion data were calculated using max and findpeak functions implemented in Matlab R2014b.

A univariate linear model analysis of covariance (ANCOVA) was conducted to determine a statistically significant difference between different study groups on the mean CV_BOLD_ values in different ROIs (GM, WM and CSF) controlling for motion parameters. This was performed by using SPSS for Windows statistical software (version 24.0; SPSS, Chicago, Illinois).

Furthermore, the effect of removal of residual motion was assessed using the additional preprocessing step of spike removal from the time-series with the AFNI 3dDespike tool using default threshold settings. After 3dDespike, the CV_BOLD_ maps were calculated and tissue-template-based mean CV_BOLD_ values were compared to the ones calculated without despiking.

#### Effect of gray matter atrophy

Structural data were analyzed with FSL-VBM, a voxel-based morphometry-style analysis [Ashburner and Friston, 2000; Good et al., 2001]. Structural images were brain-extracted using BET [Smith, 2002]. This procedure was verified with visual inspection of the extraction results. Tissue-type segmentation into GM, WM and CSF was carried out using FAST4 (55). The resulting GM partial volume images were then aligned to the Montreal Neurological Institute’s (MNI152) standard structural space template using the affine registration tool FLIRT [Jenkinson et al., 2002; Jenkinson and Smith, 2001], followed optionally by nonlinear registration using FNIRT (www.fmrib.ox.ac.uk/analysis/techrep), which uses a b-spline representation of the registration warp field [Rueckert et al., 1999].

To analyze the between-group differences on GM atrophy patterns, these resulting images were averaged to create a study-specific template, to which the native GM images were then nonlinearly re-registered. The registered partial volume images were then modulated to correct for local expansion or contraction by dividing by the Jacobian of the warp field. The modulated segmented images were then smoothed with an isotropic Gaussian kernel with a sigma of 4 mm. Finally, GM differences between different study groups were statistically tested using permutation-based nonparametric testing incorporating TFCE implemented in the FSL randomise tool with 10,000 random permutations [Smith and Nichols, 2009]. Resulting statistical maps were thresholded at p<0.05 (TFCE-corrected for familywise errors).

To analyze the effect of GM atrophy on CV_BOLD_ values GM volume in voxels was correlated subjectwise with mean CV_BOLD_ values (within GM, WM and CSF) using Spearman’s rank correlation coefficient. Scatter plots were used to visualize the results. The effect of GM atrophy was also evaluated repeating the FSL randomise analysis but using this time the GM volume as a regressor. The resulting statistic maps were spatially correlated to the ones without GM regressor using fslcc tool from FSL.

#### Receiver operating characteristic curves

We plotted receiver operating characteristic (ROC) curves to evaluate whether CV_BOLD_ could be used to separate healthy controls from patients with either AD or bvFTD, or patient groups from each other. The mean CV_BOLD_ was calculated using different ROIs: GM, WM, CSF (Fig. 2) and disease-specific templates (Fig. 3A). Area under the curve (AUC) was calculated as a measure of classification accuracy. The bootstrap approach was used to estimate the 95% confidence interval of AUC in SPSS v24.

**Fig. 2.**
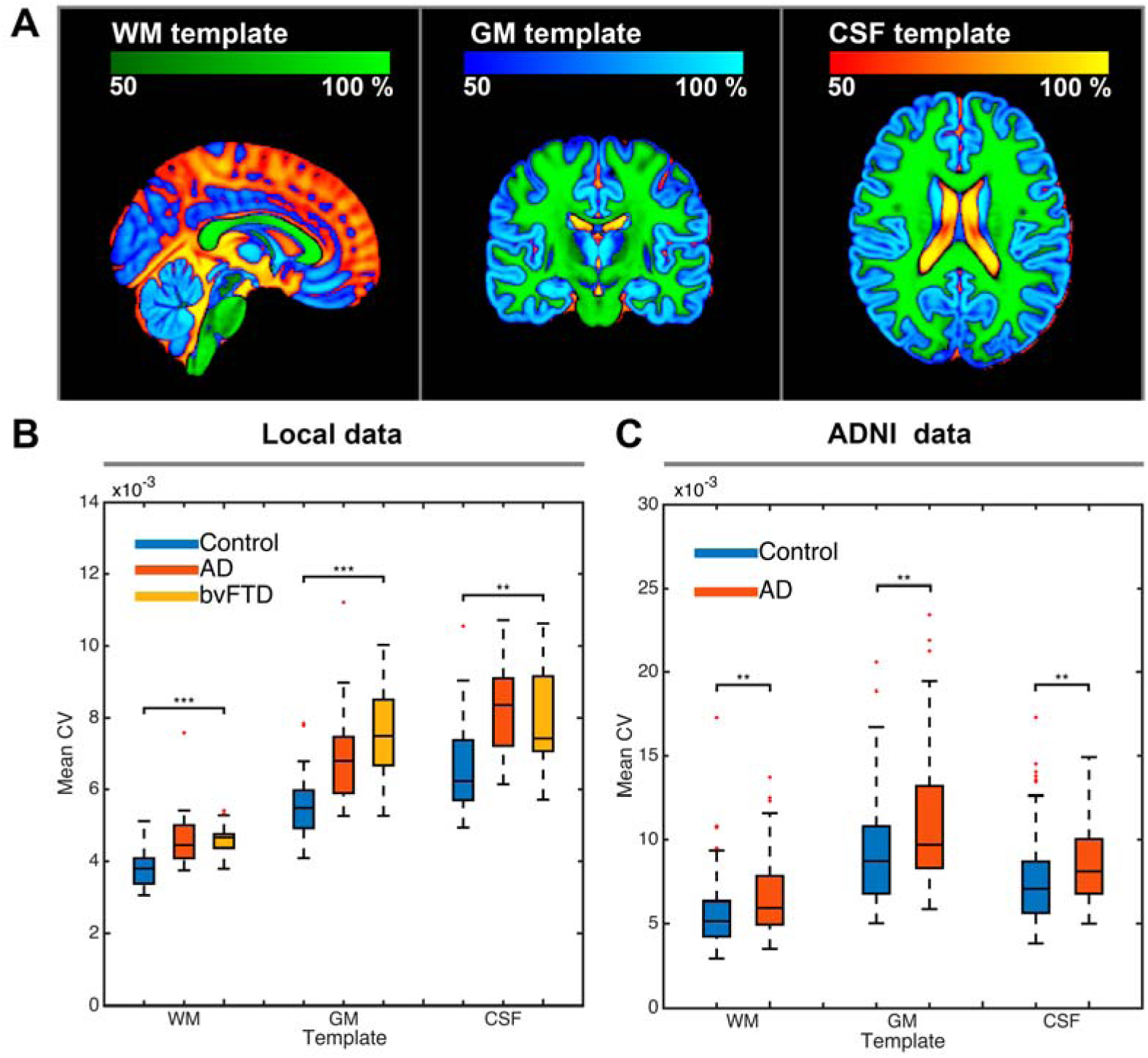
Mean CV_BOLD_ values in GM, WM and CSF. On top is the ICBM152 template used as ROI for calculating mean CV_BOLD_ values that are represented here as a boxplot with respective color encoding. Additional motion correction using despiking did not affect these results.

**Fig. 3.**
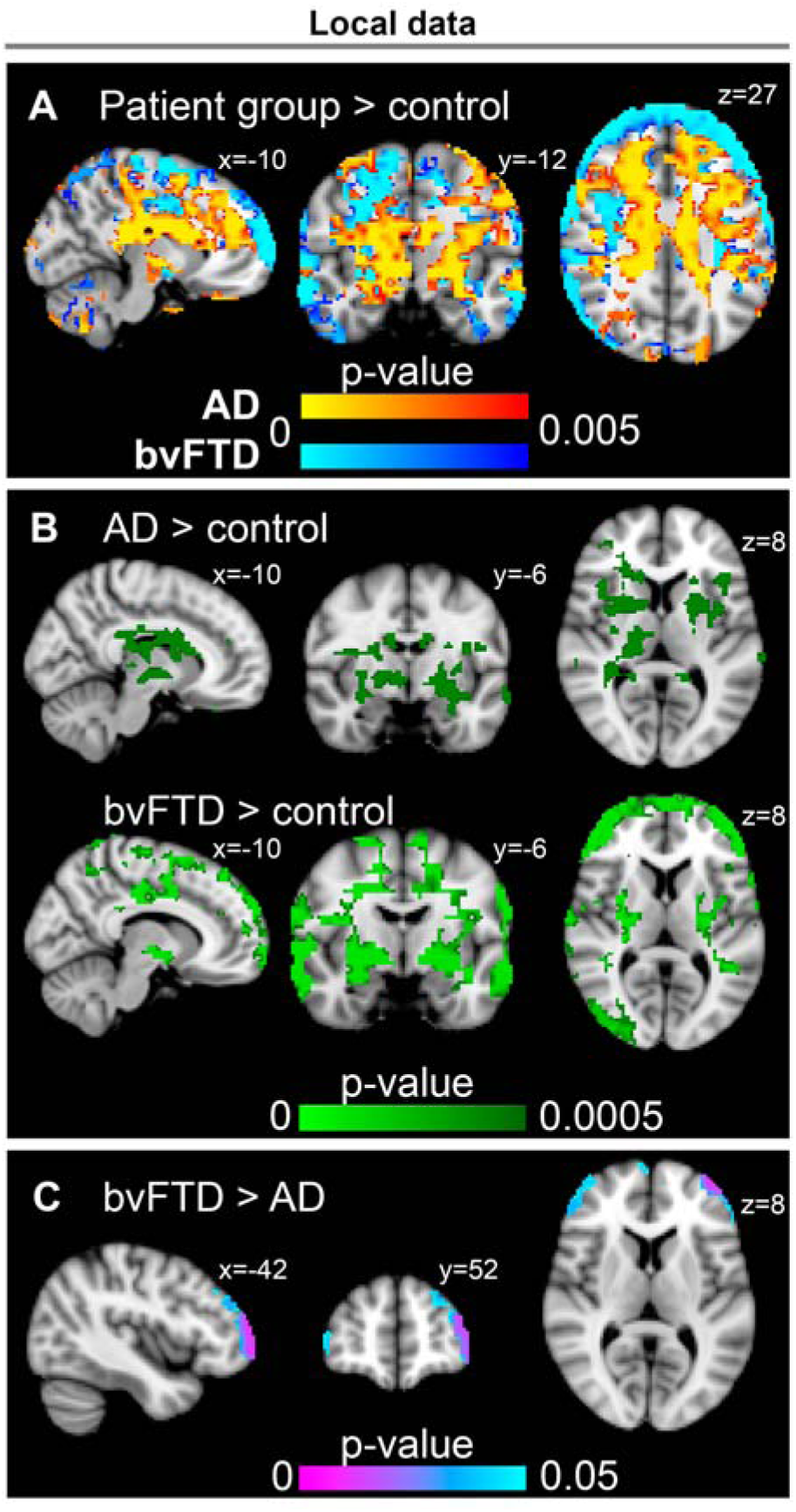
Statistically significant voxel-wise CV_BOLD_ differences between study groups and their TFCE-corrected p-values. A and B shows the AD and bvFTD results with different p-value thresholds indicated in the image. In C, the cluster of voxels is presented where CV_BOLD_ values are higher in the bvFTD group vs. AD group.

#### Statistical Analysis

Statistical analyses were performed with the SPSS and Matlab software, and p values of less than 0.05 were considered to indicate a significant difference for all analyses. Between-group differences were assessed using Kruskal-Wallis, two-tailed t and X^2^ tests, as appropriate. Subject-wise mean motion and GM volume values were correlated with the mean CV_BOLD_ values using Spearman’s rank correlation coefficient.

## Results

### Characteristics of Participants

A total of 64 healthy controls, 47 AD and 18 bvFTD patients were eligible for analysis. Demographics and clinical data are summarized in Table 1 for the local institute and in Table 2 for the ADNI data.

**Table 1.** Descriptive characteristics of the study groups in local institute data.

| Local institute data | Controls | AD | bvFTD |
| --- | --- | --- | --- |
| Participants | 24 | 17 | 18 |
| Age (yrs) | 60.0 ± 5.1 | 60.0 ± 5.4 | 60.2 ± 7.3 |
| Female | 12 (50 %) | 11 (65 %) | 9 (50 %) |
| Disease duration (yrs) | - | 2.6 ± 1.3 | 2.9 ± 1.9 |
| MMSE | 29.0 ± 1.1 * | 23.0 ± 2.6 | 24.2 ± 4.1 |
| FBI | NC | NC | 23.5 ± 4.7 |
| BDI | 3.2 ± 3.3 | NC | NC |
| Medication |  |  |  |
| Memantine | 0 | 2 | 3 |
| Acetylcholin-esterase inhibitors | 0 | 14 | 5 |
| Neuroleptic | 0 | 4 | 8 |
| Valproate | 0 | 0 | 4 |

Values represent mean±SD or N (%). AD = Alzheimer's disease. bvFTD = behavioral variant frontotemporal dementia. MMSE = Mini Mental State Examination (maximum total score is 30). FBI = Frontal Behavioral Inventory Score (maximum total score is 72). BDI = Beck's Depression Inventory (maximum total score is 63). NC = not collected. \* p < 0.05.

**Table 2.**
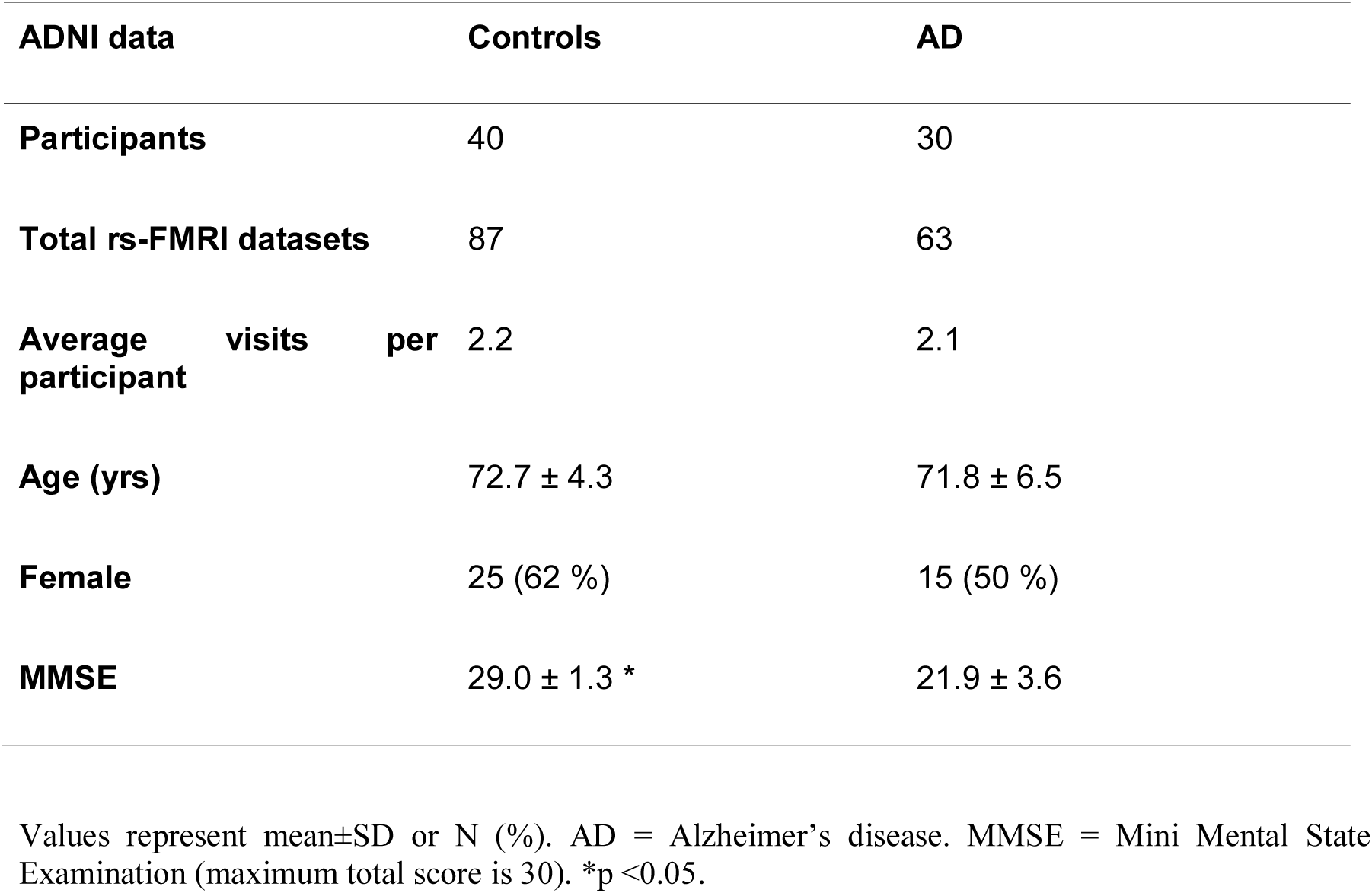
Descriptive characteristics of the study groups in ADNI data.

### CV_BOLD_ is elevated in both AD and bvFTD

Both AD and bvFTD patients showed higher CV_BOLD_ values than the controls based on visual inspection of the CV_BOLD_ maps. The group mean and examples of single-subject CV_BOLD_ maps are shown in Fig. 1.

To analyze this further, template-based ROI-analysis was conducted using ICBM152 tissue-template for WM, GM and CSF (Fig. 2). The CV_BOLD_ values on average were higher in both patient groups than in the control group (p-values ranging from 0.008 to 0.00001). This difference was confirmed using data from the ADNI study. The lowest CV_BOLD_ values were detected in the WM in all of the study groups. In the bvFTD group, mean CV_BOLD_ values were higher in WM and GM compared to both AD and the control group (Fig. 2). In the AD group, the mean CV_BOLD_ values were higher in the CSF compared to bvFTD and control group. However, the difference between bvFTD and AD groups in the large-scale ROI-analysis did not reach statistical significance (p=0.27).

### The CV_BOLD_ demonstrates disease-specific changes

More-detailed voxel-level differences of CV_BOLD_ values between study groups were analyzed using permutation-based nonparametric testing incorporating TFCE with 10,000 random permutations. This revealed distinct differences between the patient and control groups. In the AD group, significantly higher CV_BOLD_ values compared to control group were located closer to the center in periventricular WM; in GM higher CV_BOLD_ values are located in the parietal, occipital and posterior part of frontal lobes as well as in frontal pole. In the bvFTD group differences extend more to external parts of the WM and towards the frontal and temporal lobes as well as the middle occipital gyri. There are also higher CV_BOLD_ values in the cerebellum near the 4th ventricle in both diseases. In AD higher CV_BOLD_ values located more towards the horizontal fissure in posterior lobe. In order to pinpoint the most significant changes, the most statistically significant differences are illustrated with p < 0.005 and p < 0.0005 in Fig. 3 and in supplementary Fig. S1.

Statistically significant (p < 0.005) voxel-wise differences between AD patients and controls showed increased CV_BOLD_ accumulated in a circular area around the CSF ventricles centered on WM. The increased CV_BOLD_ values are found symmetrically in corpus callosum, thalamus, putamen, sagittal stratum, insula and also amygdala and anterior hippocampi areas as well as cerebellum. In the GM increased CV_BOLD_ are found in Broca’s areas, somatosensory, supplementary and sensorimotor SM1 cortices, paracingulate gyri, and also in visual V1–V3 cortices. Notably, there was no statistically significant difference between AD and control group in the parts of the DMN (posterior cingulate, angular and medial prefrontal gyri). The spatial localizations of the statistically significant changes in CV_BOLD_ values were markedly similar to the results of the ADNI data (Fig. 4).

**Fig. 4.**
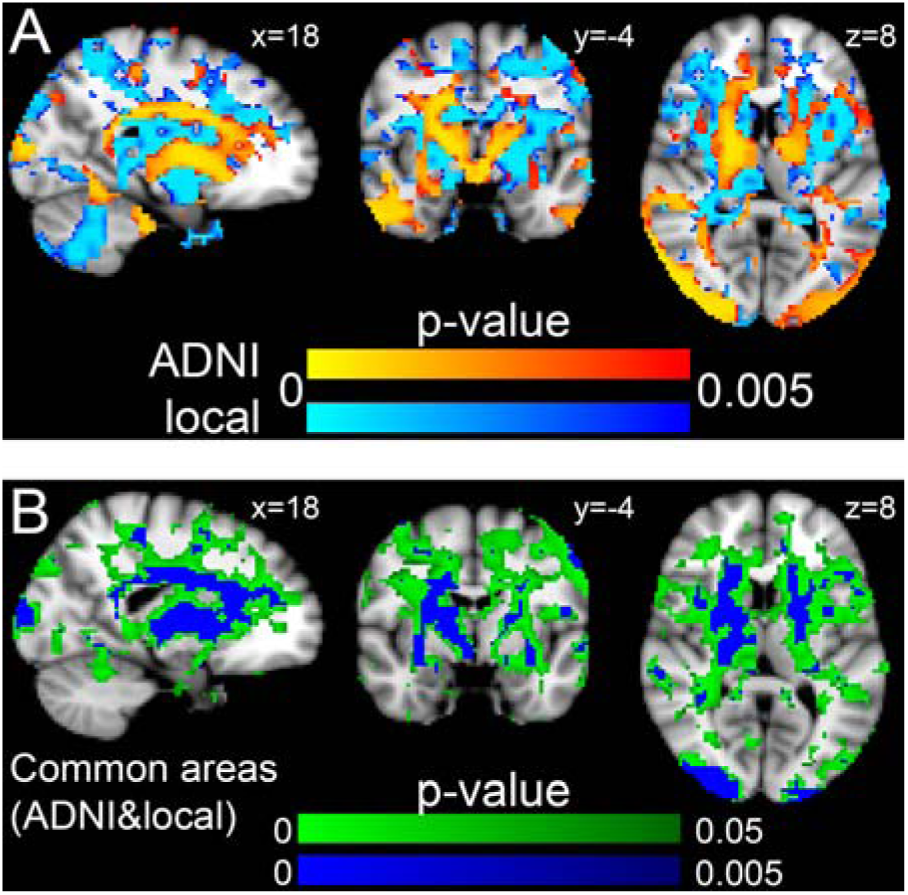
Comparison of the voxel-wise CV_BOLD_ differences between local institute and ADNI data (AD>control). B shows the common areas and mean p-value between local institute and ADNI data in two different p-values (p<0.05 and p<0.005, TFCE-corrected for familywise errors). 56 % of the statistically significant voxels (p<0.05) are shared in both data sets.

The most prominent changes in bvFTD patients showed increased CV_BOLD_ values more towards the frontal areas and lateral periventricular structures, and towards the temporal pole, premotor cortex and temporal fusiform cortex, and also in visual areas V3–V5 in lateral occipital cortex. Bilateral amygdala, putamen, insula, hippocampus, and areas in the cerebellum showed also higher CV_BOLD_ values (Fig. 3).

The statistically significant differences between the AD and bvFTD (bvFTD>AD) patients on voxel-level CV_BOLD_ were located bilaterally in anterior part of the frontal lobe (Fig. 3C).

The supplementary Tables S1–3 show the most significant group difference clusters and their anatomical labeling in the local institute data.

### CV_BOLD_ alterations are not explained by head motion or gray matter atrophy

There were no significant differences in the absolute or relative head motion parameters between any of the study groups in the local institute data (Fig. 5). In the ADNI dataset, the AD patients moved more before the motion correction (absolute: 0.20 mm for the control group and 0.30 mm for the AD group, p=0.03; relative: control 0.15 mm and AD 0.21 mm, p=0.02). Head motion did not exceed half a voxel size.

**Fig. 5.**
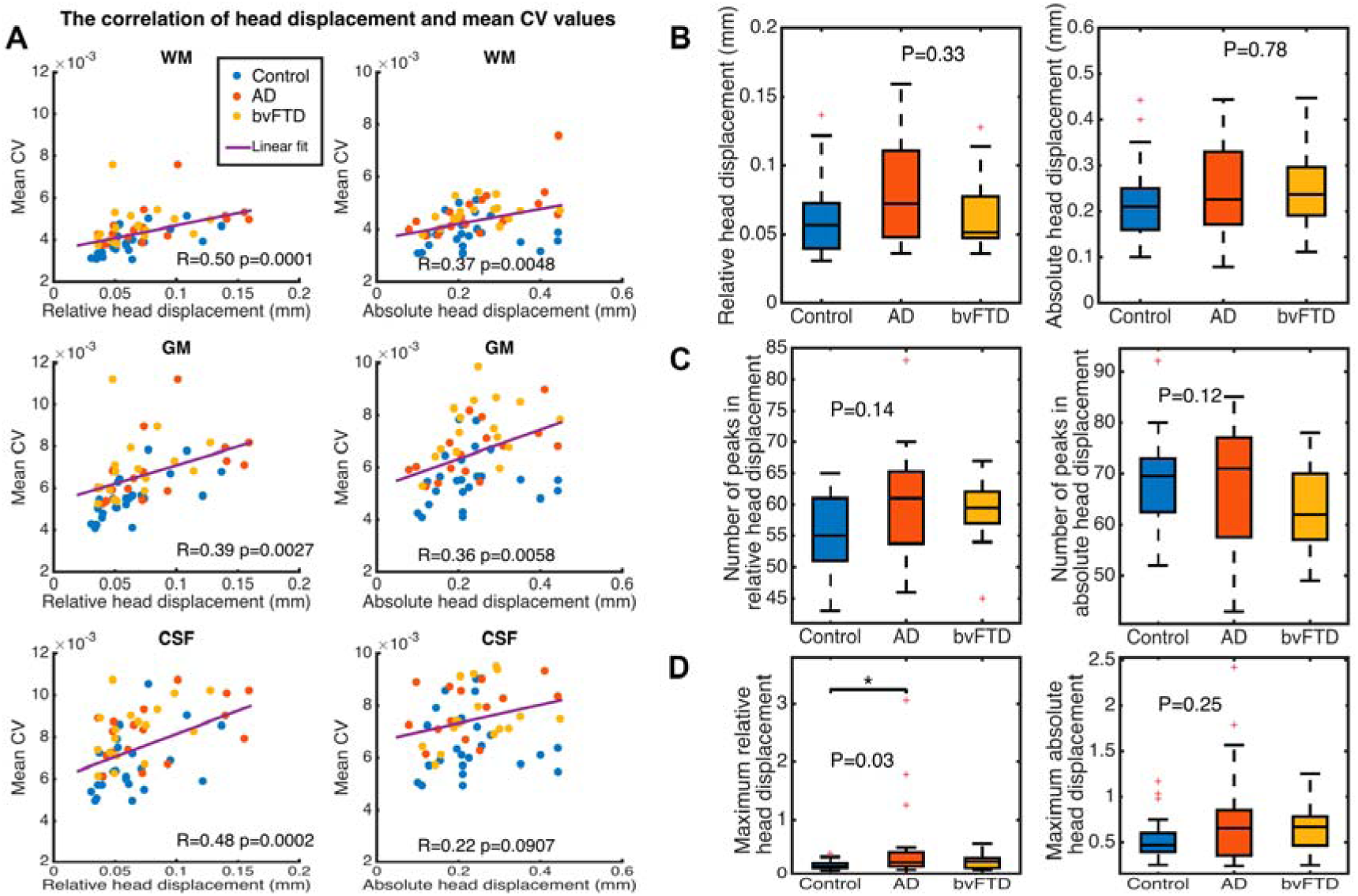
Effect of motion on CV_BOLD_ values (local institute data). Scatter plot (A) for the mean CV_BOLD_ value within template vs. head displacement (relative and absolute). Correlation between mean CV_BOLD_ and head displacement is also shown. The mean CV_BOLD_ values are higher in the patient groups (red and yellow circles) than in the control group (blue circles) with the same amount of motion both in the absolute and relative head displacement. Subject-wise absolute displacement values (in mm) were extracted, describing the amount of movement in all directions over the whole scan as a marker of gross motion. Also, relative displacement values were extracted, as a marker of motion between each EPI volume. Boxplot of the mean absolute and relative head displacement is shown in B. Differences in subject-wise mean absolute and relative motion values with CV_BOLD_ values were tested using Spearman’s rank correlation coefficient. Additionally, a number of peaks (C) and the maximum value (D) in the subject-wise motion data were evaluated.

Mean CV_BOLD_ values did correlate to motion (Fig. 5A, correlation coefficient R ranging from 0.22 to 0.50). However, there is still a highly statistically significant effect of study group on the mean CV_BOLD_ values after controlling for motion parameters, c.f. Table 3 (ANCOVA).

**Table 3.** Effect of motion.

| Template<br>(ROI) | Absolute head<br>motion |  | Relative head motion |  |
| --- | --- | --- | --- | --- |
|  | F(2,54) | Sig.<br>(p-value) | F(2,54) | Sig.<br>(p-value) |
| **WM** | 10.089 | 0.000189 | 9.584 | 0.000274 |
| **GM** | 15.006 | 0.000007 | 15.32 | 0.000005 |
| **CSF** | 7.191 | 0.002 | 6.023 | 0.004 |

Results of the ANCOVA analysis (local data). There is a statistically significant difference between different study groups on the mean $CV_{BOLD}$ values in different ROIs (GM = gray matter, WM = white matter, CSF = cerebrospinal fluid) even when controlling for absolute or relative head motion parameters.

The number of peaks in the motion signal was also analyzed as markers of sudden head movement. No statistically significant differences were found between groups and there was no correlation to the CV_BOLD_ values (data not shown, R ranging from −0.12 to 0.22, p>0.05 [0.17 to 0.45]).

We further verified the effects of motion by performing scrubbing of residual motion spikes (AFNI 3dDespike) and repeated the analysis of CV_BOLD_ group differences. The results were not affected by despiking, and there was no significant difference between mean CV_BOLD_ values calculated before and after 3dDespike (p>0.05 [0.68 – 0.98]).

The effect of GM atrophy on CV_BOLD_ values was analyzed using the local institute dataset (Fig. 6). There was no correlation between the mean CV_BOLD_ values and volume of GM (R=-0.01, p=0.91). The use of GM maps as a regressor in voxel-level analysis implementing FSL randomise resulted statistical maps that were 99 % the same as those without a regressor (spatial correlation coefficient R=0.99).

**Fig. 6.**
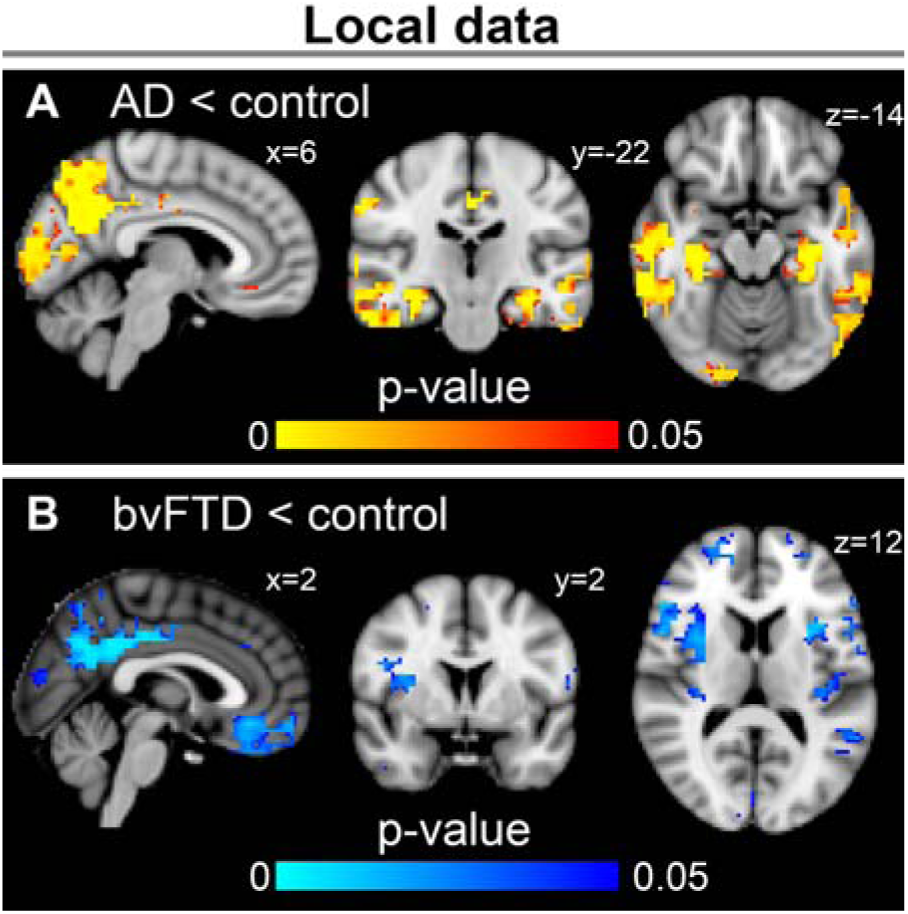
Gray matter atrophy patterns when compared to controls (local institute data). In AD, the most prominent atrophy was detected in precuneus and posterior cingulate gyrus. Significant atrophy was also detected in other temporoparietal areas. In bvFTD, atrophy was detected in posterior cingulate gyrus and also in widespread frontotemporal areas and insula.

### The accuracy of separating controls from patients with CV_BOLD_

The mean CV_BOLD_ calculated from the disease-specific templates showed excellent diagnostic accuracy. Both AD and bvFTD can be differentiated from the controls in local data with 0.96 ROC AUC values. The method also enables differentiation between AD and bvFTD, AUC being 0.806 (Fig. 7, Table 4).

**Fig. 7.**
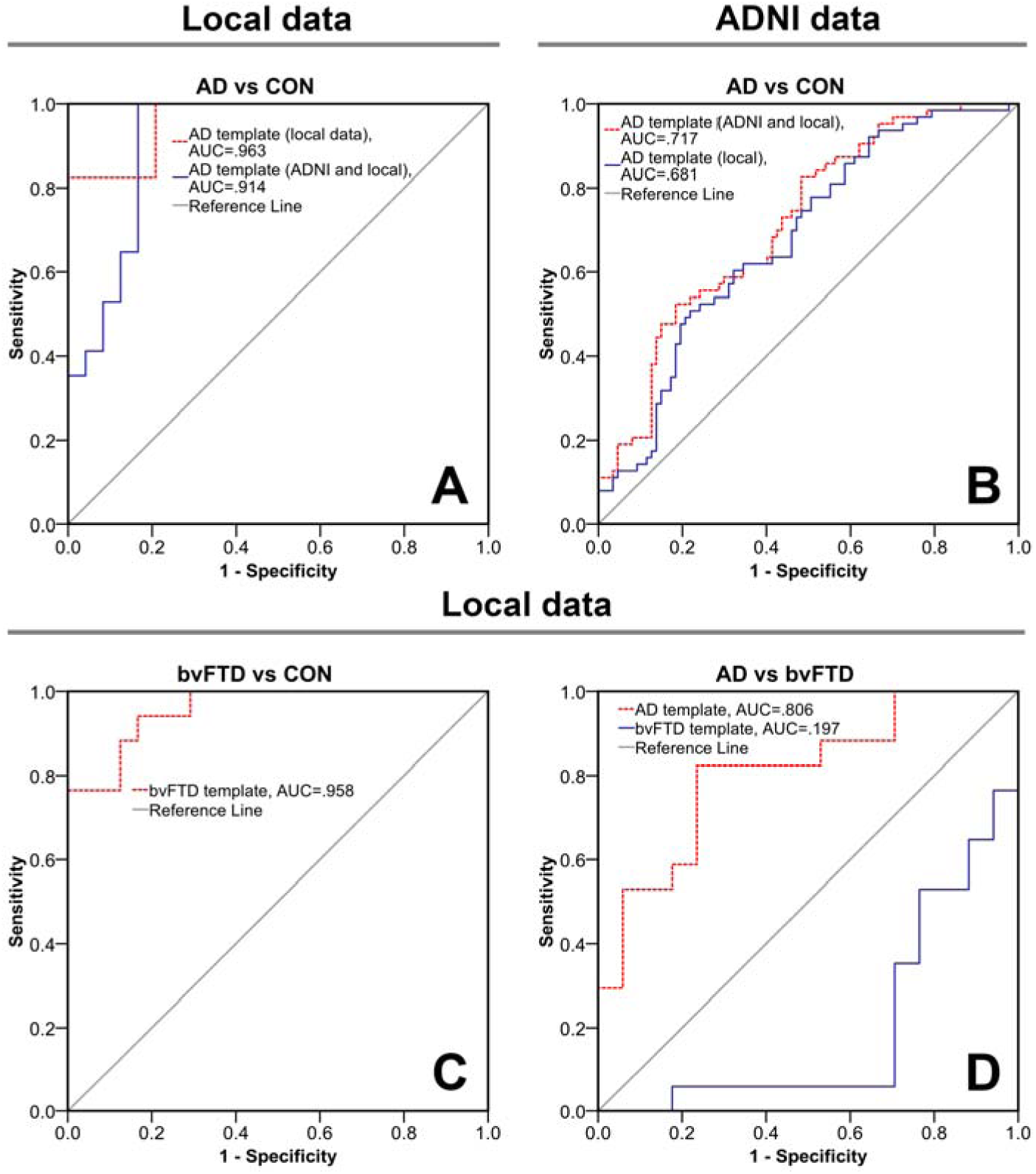
Receiver operating characteristic (ROC) curves and area under the curve values (AUC, p<0.003) for differential diagnosis based on mean CV_BOLD_ values within region-of-interest. A, ROC curve for distinguishing patients with AD from control subjects on the basis of CV_BOLD_ values within AD template as ROI. Red lines represent the template created using the most significant differences (p<0.0005) between groups in local data (Fig. 3B) and in blue the significant differences (p<0.005) common in both ADNI and local data (Fig. 4B). B, Same as in A for the ADNI data. C, ROC curve for distinguishing patients with bvFTD from control subjects on the basis of CV_BOLD_ values within the bvFTD template as ROI. The template for ROI was created using the most significant differences (p<0.0005) between groups in local data (Fig. 3B). D, ROC curve for distinguishing patients with bvFTD from those with AD on the basis of AD or bvFTD template. Confidence intervals are shown in Table 4.

**Table 4.**
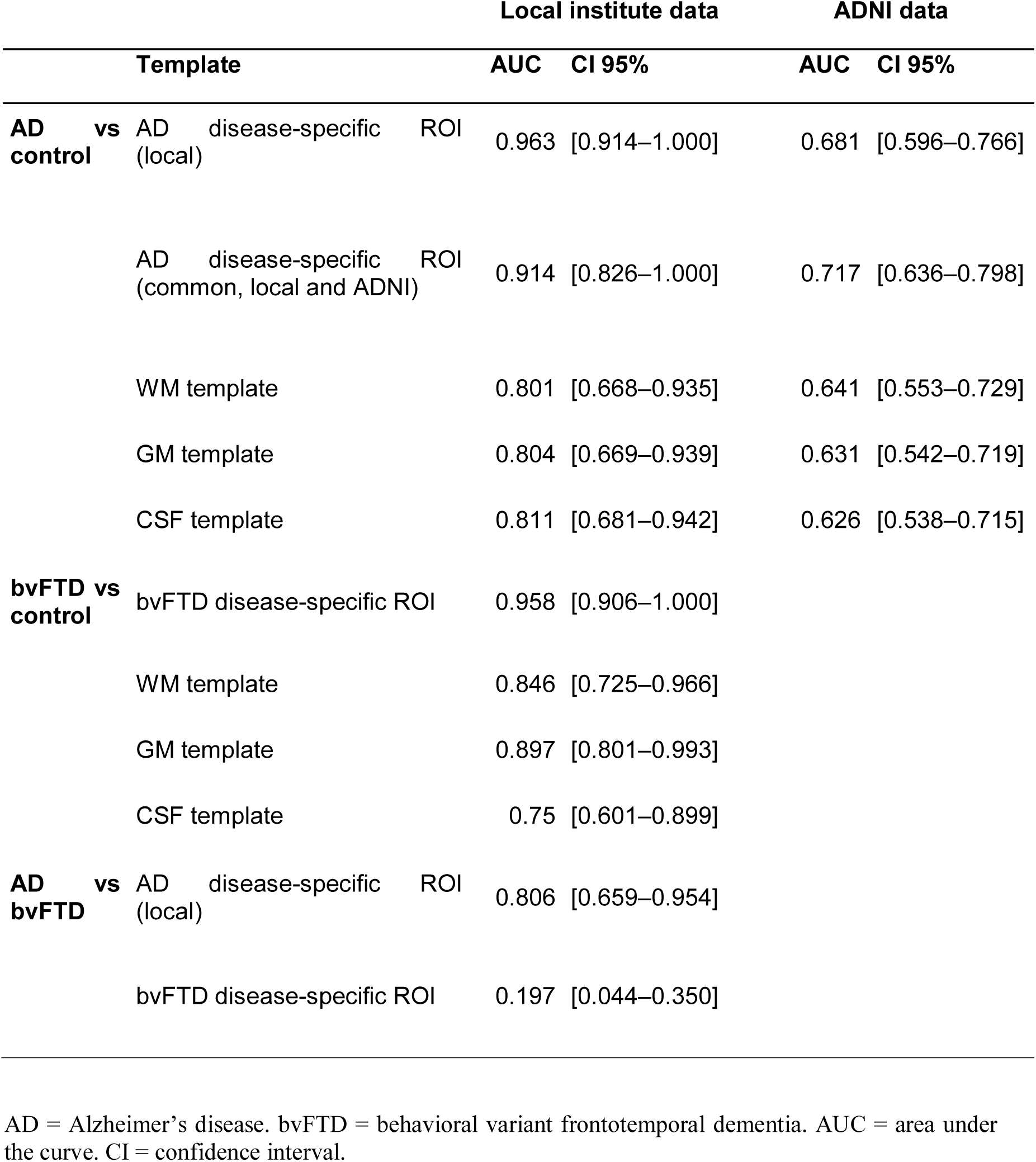
Results of the ROC analysis.

## Discussion

The goal of this study was to investigate if physiological signal contributions of BOLD data measured using CV_BOLD_ are altered in different types of dementia. We found that CV_BOLD_ is markedly increased in both AD and bvFTD compared to age-matched controls (p<0.0005), and that the CV_BOLD_ changes are motion and GM atrophy independent and therefore presumably intrinsic physiological changes. Localizations of the CV_BOLD_ alterations are somewhat disease-specific. However in both diseases, the most profound changes in the CV_BOLD_ involve areas surrounding CSF, extending to WM, basal ganglia and multiple cortical structures. Suiting the disease pathology of the bvFTD, the most significant differences in CV_BOLD_ comparison to AD were detected in frontolateral GM areas. Mean CV_BOLD_ in the disease-specific templates was able to discern AD patients from controls with the receiver operating characteristic AUC values of 0.963 and bvFTD patients from controls with an AUC value of 0.958. The AD and bvFTD groups were separated from each other with an AUC value of 0.806.

Garrett et al. has suggested that it would be beneficial to broaden the analysis of BOLD signal to variability, as it seems to be more than just noise [Garrett et al., 2010]. BOLD signal variability has been measured with standard deviation of BOLD signal (SD_BOLD_) [McIntosh et al., 2010; Wang et al., 2008]. SDBOLD has been found to reflect brain status, and that is also related to aging, pathology, cognitive skills and AD [Garrett et al., 2010; Garrett et al., 2011; Garrett et al., 2013; Garrett et al., 2015; Grady and Garrett, 2014; Grady and Garrett, 2018; Guitart-Masip et al., 2016; Scarapicchia et al., 2018]. Recently, CV_BOLD_ has been used to analyze physiological noise characteristics of BOLD data [Jahanian et al., 2014]. CV_BOLD_ has been shown to correlate with cerebral blood volume and cerebral blood flow [Khalil et al., 2017]. The theoretical advantage of CV_bold_ over SD_bold_ is that any intensity level changes are normalized to the average BOLD signal level of the voxel minimizing the effects of local things like susceptibility alterations. While BOLD signal has conventionally been related to GM, recent studies have shown increasing evidence of WM contribution to BOLD signal, as well as that there are disease-related changes in it [Ding et al., 2018; Gawryluk et al., 2014; Özbay et al., 2018; Peer et al., 2017]. Using a similar approach as CV_BOLD_ Makedonov et al. has shown increased physiological fluctuations in WM in AD [Makedonov et al., 2016] and in small vessel disease [Makedonov et al., 2013]. Makedonov et al. has suggested that the BOLD signal variation reflects end-arteriole intracranial pulsatility effects [Makedonov et al., 2013].

AD is usually been considered as a GM disease due to the distribution of hallmark pathological changes such as abnormal extracellular aggregates of amyloid-beta (Aβ) protein and intracellular neurofibrillary tangles of hyperphosphorylated tau protein [Selkoe, 1991; Ryan et al., 2015]. However, the pathogenesis of AD is still controversial. AD is also linked to loss of synapsis and myelin, mitochondrial dysfunction, oxidative stress, metabolic disorders, neuroinflammation and loss of cholinergic and other neurons [Holtzman et al., 2011; Audano et al., 2018; Bartzokis, 2011; Mark et al., 2015; Olsson et al., 2016; Tracy and Gan, 2018; Wang et al., 2016]. Patients with clinically diagnosed AD have commonly mixed AD and cerebrovascular disease pathology, and there has been much interest in the interactions between these diseases [Toledo et al., 2013]. The cardiovascular and respiratory pulsations drive the glymphatic clearance of the brain. Glymphatic failure has been strongly linked to neurodegenerative diseases [Iliff et al., 2012; Iliff et al., 2013; Iliff et al., 2014; Iliff et al., 2015; Kiviniemi et al., 2016; Kress et al., 2014; de Leon et al., 2017; Louveau et al., 2016; Peng et al., 2016; Plog et al., 2015; Snyder et al., 2015; Tarasoff-Conway et al., 2015]. B-amyloid has been found to increase in the periventricular WM already after one night sleep deprivation, which suggests that absence of nighttime glymphatic brain clearance surge may predispose to amyloid deposition and neuronal degeneration [Shokri-Kojori et al., 2018; Xie et al., 2013].

Cerebral small vessel disease is increasingly linked to cognitive decline and dementia [Bos et al., 2018]. On T2-weighted MRI, white matter disease (WMD) is represented as white matter hyperintensities (WMH), which are thought to reflect demyelination and axonal loss. WMH is a good predictor of AD incidence and early AD patients with micro- and macro-structural abnormalities in the white matter have higher risk of disease progression [Brickman et al., 2012; Brickman, 2013; Brickman et al., 2015; Sjöbeck et al., 2006; Tosto et al., 2014]. Magnetic susceptibility differences has been associated with tau pathology and increased staining for reactive microglia and astrocytes [Acosta-Cabronero et al., 2013; O’Callaghan et al., 2017].

In the present study, analysis of CV_BOLD_ was not limited to WM as in previous study by [Makedonov et al., 2016]. However, CV_BOLD_ was altered in AD most dominantly in the periventricular WM structures like corpus callosum. A decrease of vessel density in the periventricular region has been observed in AD [Brown et al., 2009]. Demyelinated lesions tend to distribute within the areas with relatively low cerebral blood flow, which are usually found in profound, periventricular WM. Our results add further proof into the vascular abnormalities by showing changes in CV_BOLD_ within the WM.

We also analyzed CV_BOLD_ in different type of dementia (bvFTD). The neuropathology associated with bvFTD is heterogeneous, and at the moment there is no clear relationship between the clinical phenotype and the underlying pathogenesis. One of the most consistent pathological finding of bvFTD is the relatively selective atrophy of the frontal and temporal lobes. In most cases post-mortem immunohistopathology shows abnormal protein inclusions in neural and glial cells. Based on immunohistochemical staining there are multiple subtypes of bvFTD (major subtypes being tau and TDP) [Mackenzie et al., 2010]. bvFTD has been found to be associated with progressive degeneration of the SLN resting-state network [Lee et al., 2014; Seeley et al., 2009; Seeley et al., 2012]. SLN is a resting-state network that includes the anterior cingulate and frontoinsular cortex, amygdala, striatum, and medial thalamus. These region bridges the frontal lobes and limbic system, and it has been proposed to represent the emotional significance of internal and external stimuli and coordinate contextualized viscero-autonomic, cognitive, and behavioural responses [Lee et al., 2014; Seeley et al., 2007; Whitwell and Josephs, 2012]. The other resting-state networks have received much less interest, and there have been more controversies between results [Farb et al., 2013; Filippi et al., 2013; Hafkemeijer et al., 2012; Rytty et al., 2013; Tuovinen et al., 2017].

In areas that are part of the SLN, CV_BOLD_ is higher in bvFTD, compared to control group. Interestingly, the regions of the DMN were not found to have higher CV_BOLD_ values in the AD group. Recent study showed that bvFTD patients displayed more fixations to the eyes of the emotional faces, compared to controls [Hutchings et al., 2018]. Regions associated with fixations to the eyes included the left inferior frontal gyrus, right cerebellum and middle temporal gyrus; in this study these areas were found to have higher CV_BOLD_ values. Both of the diseases seem to affect the CV_BOLD_ measured from the cerebellum. Cerebellar atrophy have been found in both AD and bvFTD [Gellersen et al., 2017].

In the present study the significant difference between the AD and bvFTD was shown to be in bilateral frontal poles. The known neuropsychological differences between AD and bvFTD can be in part explained by the CV_BOLD_ differences between the conditions in the frontopolar areas. These areas are known have a role in resolving indeterminate relations in un-certain situations and intensity of emotions [Goel et al., 2009]. Left ventral prefrontal cortex is involved in active and strategic operation of the mnemonic representation and retrieval success for words [Iidaka et al., 2000]. Study by Wong et al. contrasted prefrontal cortex atrophy with episodic memory dysfunction in AD and bvFTD [Wong et al., 2014]. Episodic memory deficits are underpinned by divergent prefrontal mechanisms: left side frontal pole for AD and right side for bvFTD, similar to our results. Our results indicate that the most marked alterations in bvFTD occur in the right lateral areas of frontal poles with increased CV_BOLD_. A study comparing AD and bvFTD revealed non-atrophy related perfusion deficits in frontal areas in accordance with our results [Du et al., 2006].

When compared to bvFTD, CV_BOLD_ changes in AD occur more on the basal, periventricular areas where cardiovascular pulsations dominate in physiological studies [Kiviniemi et al., 2016]. The bvFTD changes are occurring more towards the frontal cortical edges of the brain, which was recently shown to be connected to respiratory brain pulsations of the glymphatic system [Kiviniemi et al., 2016]. Interestingly, arterial hypertension and other vascular risk factors are known risk factors for AD, but not for bvFTD [Baborie et al., 2011; Baborie et al., 2012; De Reuck, 2012; De Reuck et al., 2012a; De Reuck et al., 2012b; Snyder et al., 2015].

The changes in CV_BOLD_ in present study are not explained by difference in age, gender, motion or GM atrophy. There were no statistically significant differences in age, gender or motion parameters between different study groups in the local institute data. As expected, there was a positive correlation with CV_BOLD_ values and motion parameters in all groups alike. However, the patient groups had increased CV_BOLD_ values with the same amount of motion. The effect of motion was also evaluated using an ANCOVA, where differences between groups prevailed as statistically significant after motion parameters were used as covariates. Also, the effect of sudden motion “peaks” was evaluated and this did not explain the group differences in the CV_BOLD_ values. Furthermore, removal of residual motion spikes by despiking had no significant effect on the CV_BOLD_ results. GM atrophy patterns were in line with previous literature [Du et al., 2006; Hartikainen et al., 2012; Tartaglia et al., 2011; Whitwell and Josephs, 2012]. CV_BOLD_ values and GM volume had no correlation. The use of GM maps, as a regressor in voxel-level analysis did not affect the results.

The previous literature and our results suggest that the changes in CV_BOLD_ are not only due to motion, but rather the changes may be due to yet unknown intrinsic properties of the degenerated brain tissue. ADNI data may be more sensitive to hemodynamically coupled BOLD signal changes due to the higher magnetic field strength (3 T) than in local institute data (1.5 T) [van der Zwaag et al., 2009]. However, the increased motion in ADNI data may partly mask the CV_BOLD_ differences between groups. Furthermore, the local institute data is also nearly two times faster in sampling rate (TR 1.8 vs. 3 seconds). This may partially increase the sensitivity to physiological pulsation noise due to somewhat reduced aliasing with faster TR [Kiviniemi et al., 2005; Smith et al., 2007]. Furthermore, as the differences predominate in CSF and WM structures, the source of T2*-weighted GRE EPI signal's CV_BOLD_ alterations in AD and bvFTD are most likely due to physiological brain pulsations rather than secondary hemodynamic changes to neuronal activity. Both local institute and ADNI data still suffer from cardiorespiratory signal aliasing and cannot pinpoint the physiological origin of the changes in CV_BOLD_. In future studies, one could investigate the source of altered CV_BOLD_ in neurodegenerative diseases with ultra-fast, critically sampled multimodal neuroimaging data. Such data can differentiate cardiorespiratory pulsations and bring forth a mechanistic explanation to the detected alteration in CV_BOLD_.

These results should be verified with larger sample sizes and the relationship between CV_BOLD_ and clinical parameters, including precise co-analysis of structural images, should be evaluated. Previous studies have employed a number of different variations of ‘BOLD variability’ measures, with different methodologies used by different groups. There have not been comparative studies of different methodologies in this field. However, these results are inline with previous literature and together with previous studies, our findings suggest that analysis of physiological pulsations using BOLD signal variability may provide useful information in the context of neurodegenerative diseases.

### Conclusions

There are disease specific alterations in CV_BOLD._ In AD these alterations was confirmed in two different datasets and in different imaging setups (1.5 T and 3 T). CV_BOLD_ changes are motion and GM atrophy independent and therefore presumably intrinsic physiological changes. Together with previous studies, our findings suggest that analysis of physiological pulsations may provide useful information in the context of neurodegenerative diseases.

## Acknowledgements

Data collection and sharing for this project was funded by the Alzheimer's Disease Neuroimaging Initiative (ADNI) (National Institutes of Health Grant U01 AG024904) and DOD ADNI (Department of Defense award number W81XWH-12-2-0012). ADNI is funded by the National Institute on Aging, the National Institute of Biomedical Imaging and Bioengineering, and through generous contributions from the following: AbbVie, Alzheimer’s Association; Alzheimer’s Drug Discovery Foundation; Araclon Biotech; BioClinica, Inc.; Biogen; Bristol-Myers Squibb Company; CereSpir, Inc.; Cogstate; Eisai Inc.; Elan Pharmaceuticals, Inc.; Eli Lilly and Company; EuroImmun; F. Hoffmann-La Roche Ltd and its affiliated company Genentech, Inc.; Fujirebio; GE Healthcare; IXICO Ltd.; Janssen Alzheimer Immunotherapy Research & Development, LLC.; Johnson & Johnson Pharmaceutical Research & Development LLC.; Lumosity; Lundbeck; Merck & Co., Inc.; Meso Scale Diagnostics, LLC.; NeuroRx Research; Neurotrack Technologies; Novartis Pharmaceuticals Corporation; Pfizer Inc.; Piramal Imaging; Servier; Takeda Pharmaceutical Company; and Transition Therapeutics. The Canadian Institutes of Health Research is providing funds to support ADNI clinical sites in Canada. Private sector contributions are facilitated by the Foundation for the National Institutes of Health (www.fnih.org). The grantee organization is the Northern California Institute for Research and Education, and the study is coordinated by the Alzheimer’s Therapeutic Research Institute at the University of Southern California. ADNI data are disseminated by the Laboratory for Neuro Imaging at the University of Southern California.

